## Supplementary material for "Genetic compatibility and ecological connectivity drive the dissemination of antibiotic resistance genes": Supplemenary Fig. 1-13, Supplementary Table 1

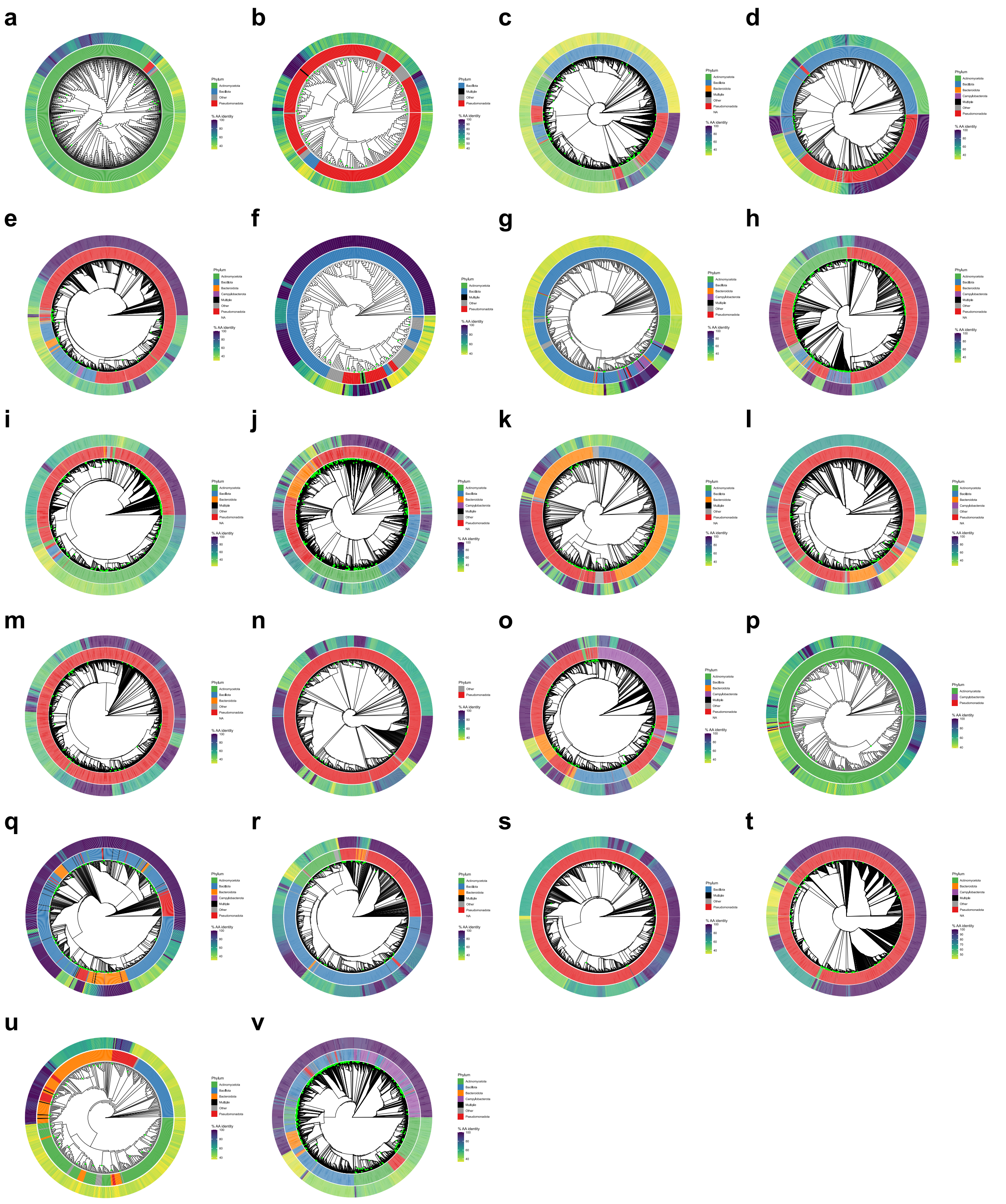

**Supplementary Fig. 1 Phylogenetic tree created from unique protein sequences predicted by fARGene, using each of the 22 available gene models.** Predicted horizontally transferred ARGs are marked as green dots in the tree. The inner circle shows the phylum of the hostto which the bacteria represented by each leaf belong, and the outer circle shows the similarity (% amino acid identity) between each protein and its closest homolog found in the CARD database. **a** Aminoglycoside model A (AAC(2')), **b** Aminoglycoside model B (AAC(3)), **c** Aminoglycoside model C (AAC(3)), **d** Aminoglycoside model D (AAC(6')), **e** Aminoglycoside model E (AAC(6')), **f** Aminoglycoside model F (AAC(6')), **g** Aminoglycoside model G (APH(2'')), **h** Aminoglycoside model H (APH(3')+APH(3'')), **i** Aminoglycoside model I (APH(6)+APH(3')), **j** Class A beta-lactamases, **k** Class B1/B2 beta-lactamases, **l** Class B3 beta-lactamases, **m** Class C beta-lactamases, **n** Class D1 beta-lactamases, **o** Class D2 beta-lactamases, **p** Erm type A, **q** Erm type F, **r** Mph, **s** Qnr, **t** Tetracycline efflux pumps, **u** Tetracycline inactivation enzymes, **v** Tetracycline ribosomal protection genes (RPGs).

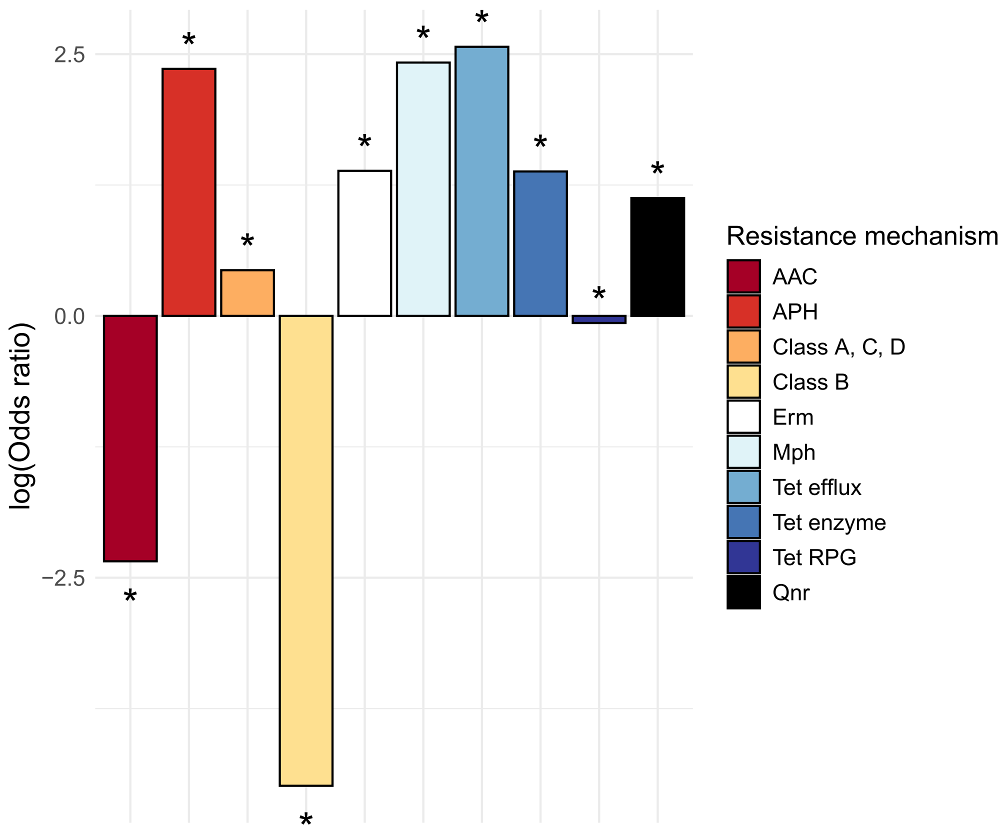


**Supplementary Fig. 2** Enrichment analysis of predicted antibiotic resistance genes involved in identified horizontal transfers. The ratios and their significance were calculated using Fisher's exact test, an asterisk is used to denote significant results (*p*<0.01).

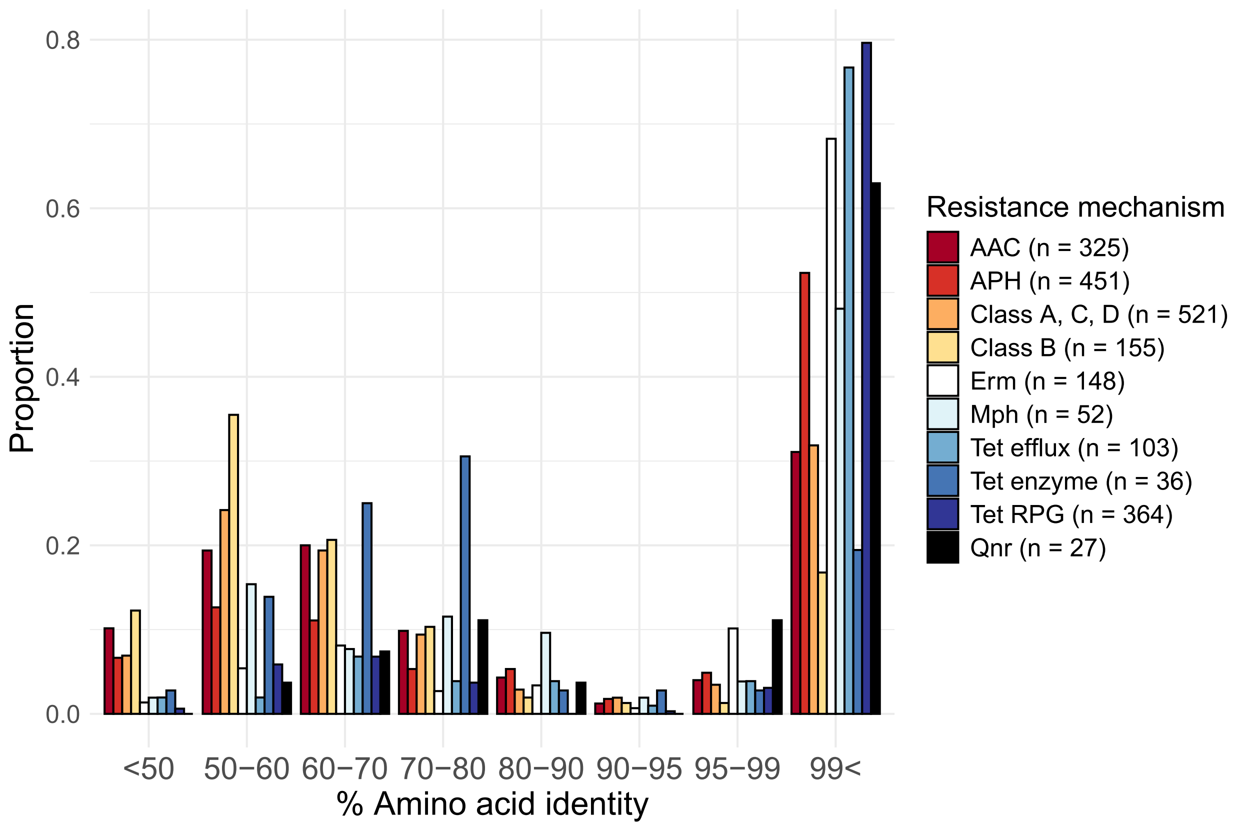


**Supplementary Fig. 3** Maximum similarity (% amino acid identity) observed between the ARGs carried by species from different taxonomic orders in the observed horizontal transfers. Here, each leaf or node in the tree where a transfer was observed is counted only once. The observations are divided based on the resistance mechanism that the transferred gene encodes.

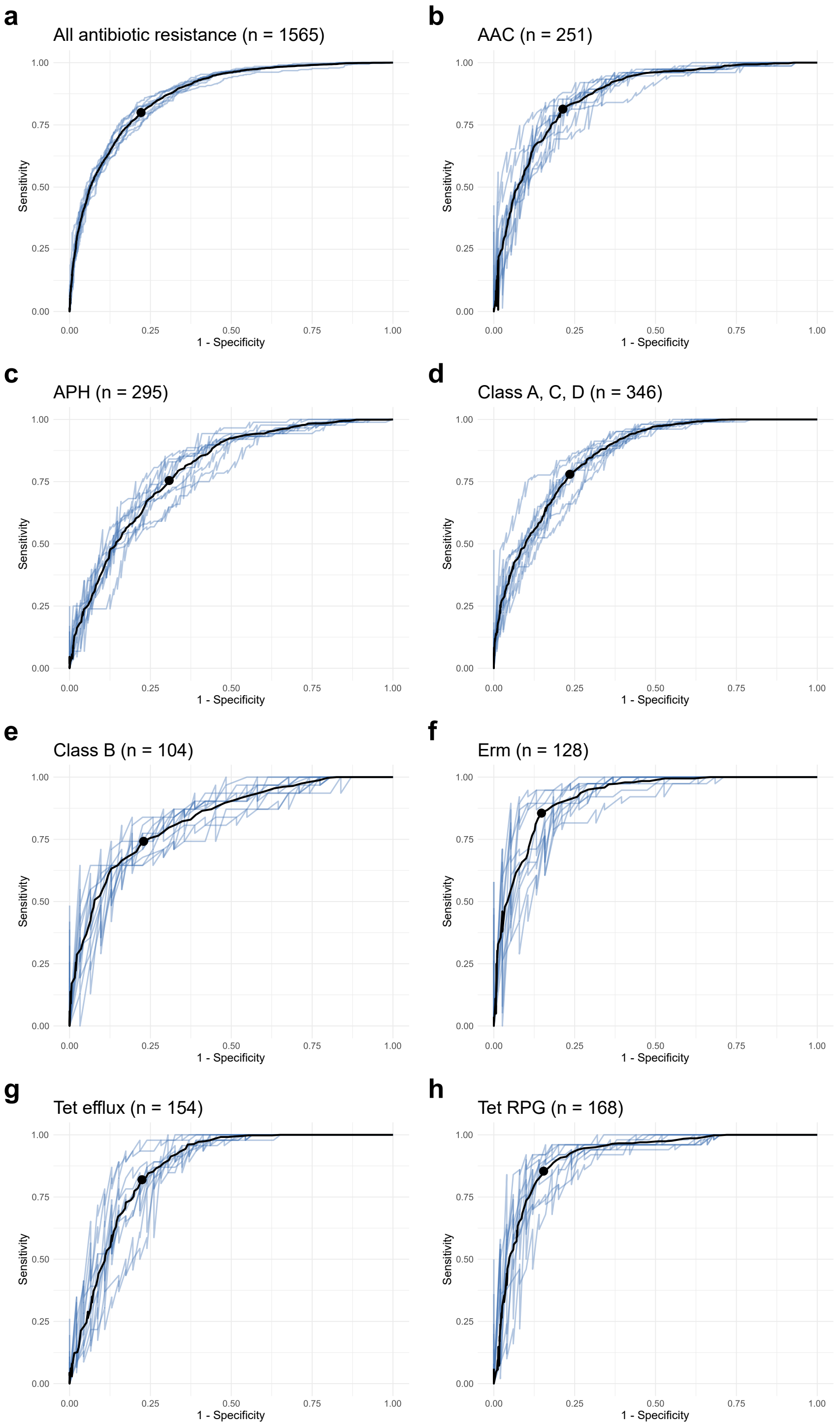


**Supplementary Fig. 4 Reciever operating characteristic curve produced from predictions on test data by random forest classifiers.** The models were trained on horizontal transfers representing **a** all included resistance mechanisms, **b** AAC aminoglycoside acetyltransferases, **c** APH aminoglycoside phosphotransferases, **d** Class A, C, D beta-lactamases, **e** Class B beta-lactamases, **f** Erm 23S rRNA methyltransferases, **g** Tetracycline efflux pumps, **h** Tetracycline ribosomal protection genes (RPG). A point is placed on each curve representing the observed optimal performance. The number of observed transfers making up the training + test data for each model is included in the titles.

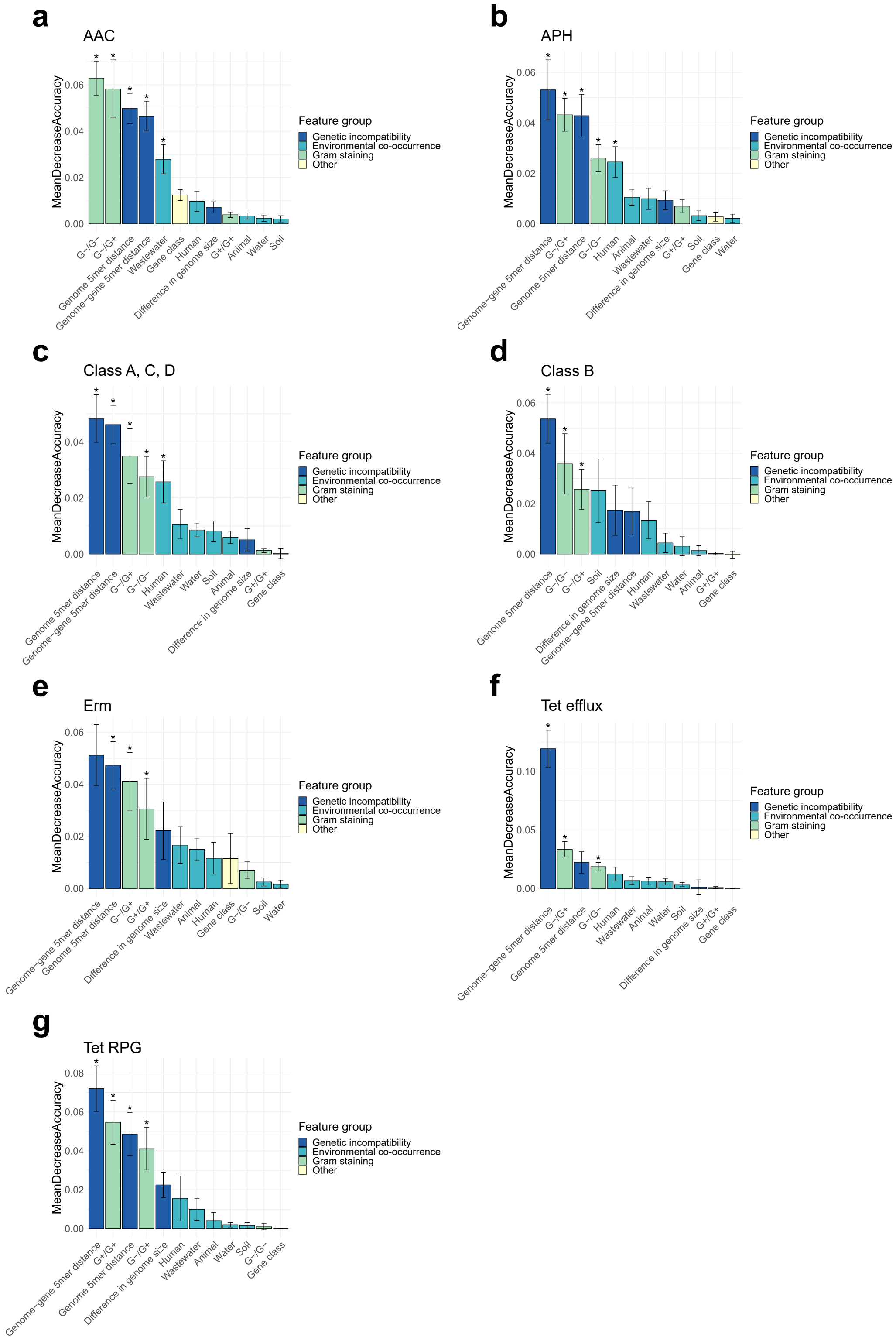


**Supplementary Fig. 5 Permutation importance of the features included in the resistance mechanisms-specific random forest classifier**. Features are ordered according to their overall contribution to the accuracy of the classifier (MeanDecreaseAccuracy). **a** AAC aminoglycoside acetyltransferases. **b** APH aminoglycoside phosphotransferases. **c** Class A, C, D beta-lactamases. **d** Class B beta-lactamases. **e** Erm 23S rRNA methyltransferases. **f** Tetracycline efflux pumps. **g** Tetracycline ribosomal protection genes (RPG).

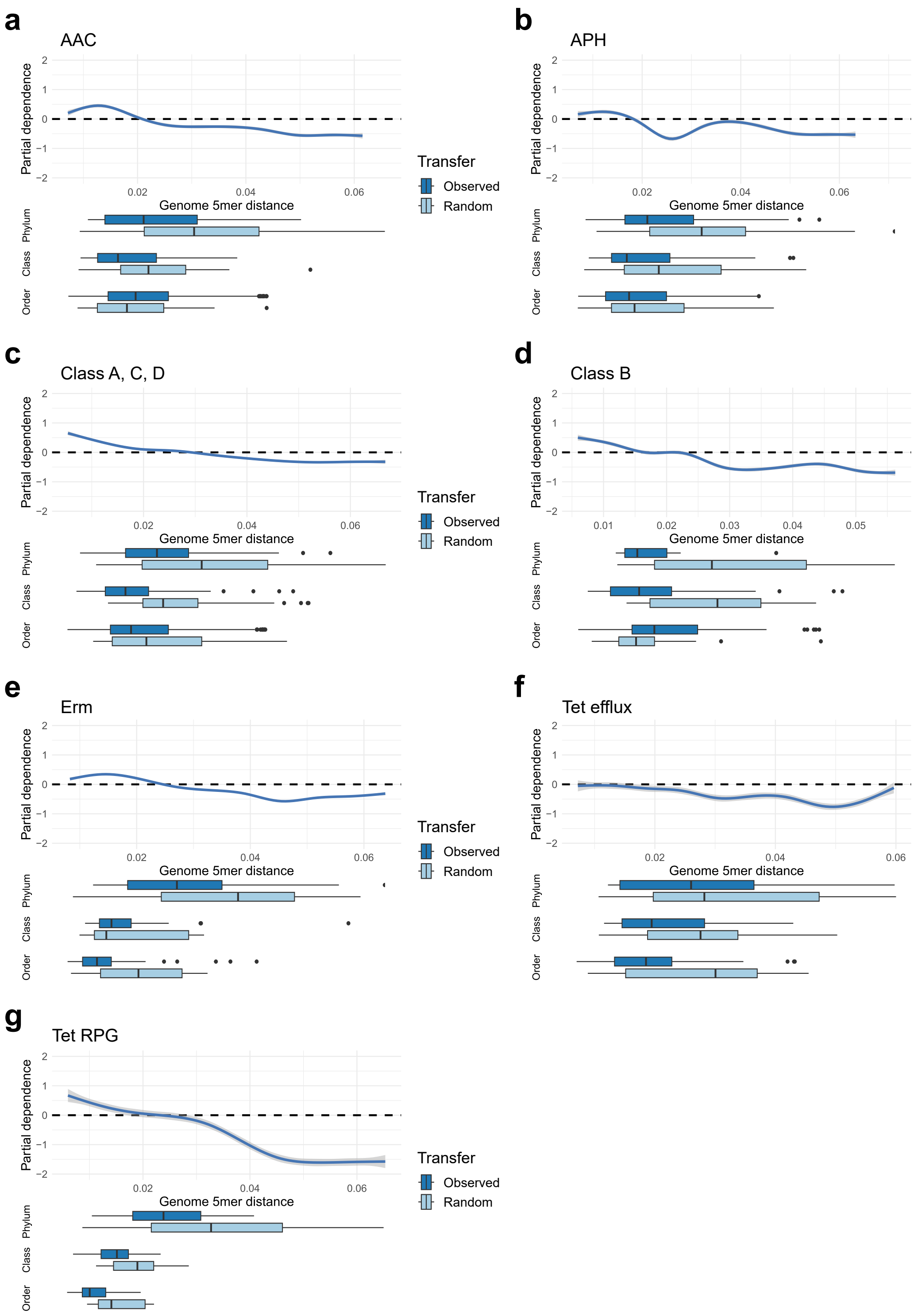


**Supplementary Fig. 6 Relative contribution of the genome 5mer distance to the classification of horizontally spread ARGs for the mechanism-specific random forest models.** The distribution of values observed *the obaserved and the randomized transfers* at different taxonomic levels is visualized as boxplots below the main graph. **a** AAC aminoglycoside acetyltransferases. **b** APH aminoglycoside phosphotransferases. **c** Class A, C, D beta-lactamases. **d** Class B beta-lactamases. **e** Erm 23S rRNA methyltransferases. **f** Tetracycline efflux pumps. **g** Tetracycline ribosomal protection genes (RPG).

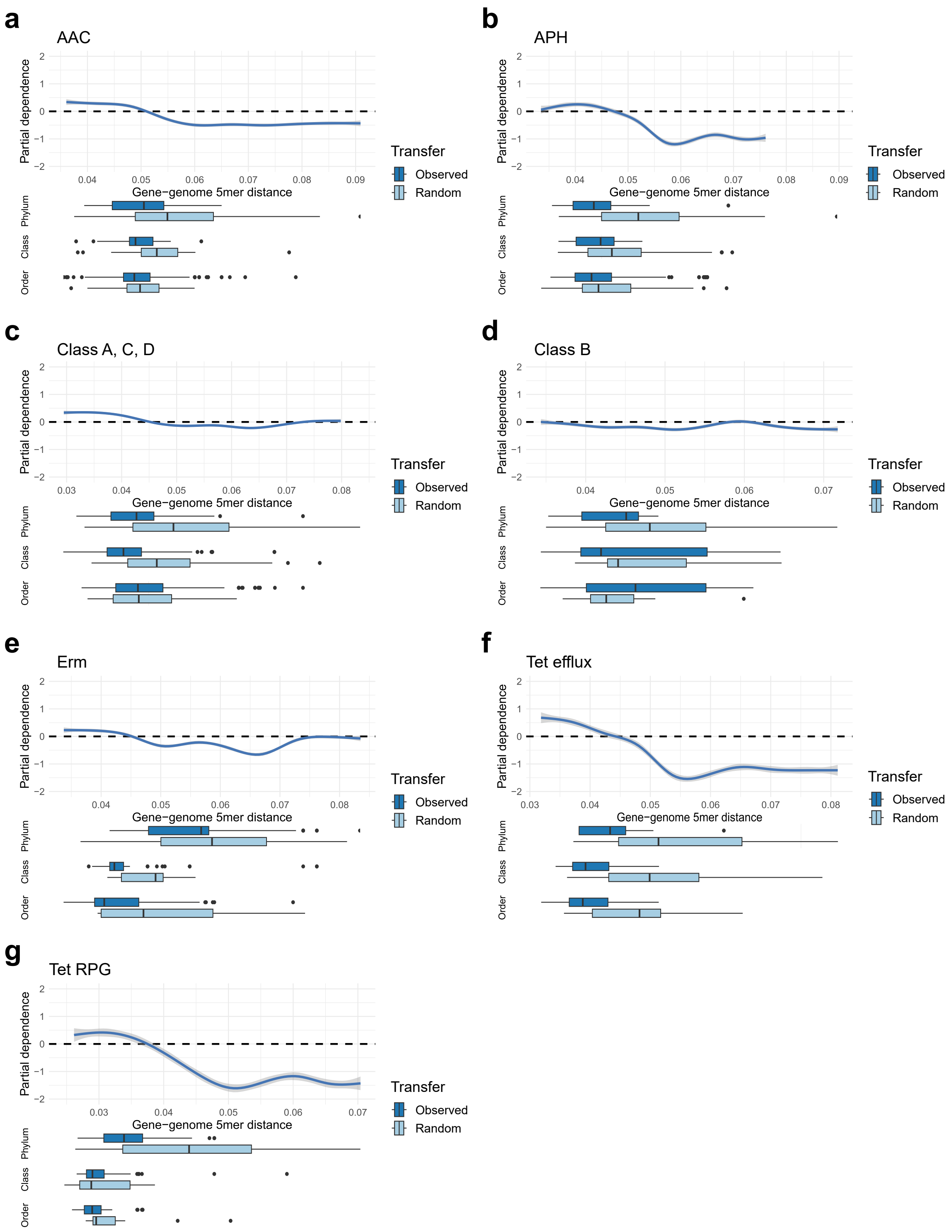


**Supplementary Fig. 7 Relative contribution of the gene-genome 5mer distance to the classification of horizontally spread ARGs for the mechanism-specific random forest models.** The distribution of values seen for the observed and the randomlizeed transfers at different taxonomic levels is visualized as boxplots below the main graph. **a** AAC aminoglycoside acetyltransferases. **b** APH aminoglycoside phosphotransferases. **c** Class A, C, D beta-lactamases. **d** Class B beta-lactamases. **e** Erm 23S rRNA methyltransferases. **f** Tetracycline efflux pumps. **g** Tetracycline ribosomal protection genes (RPG).

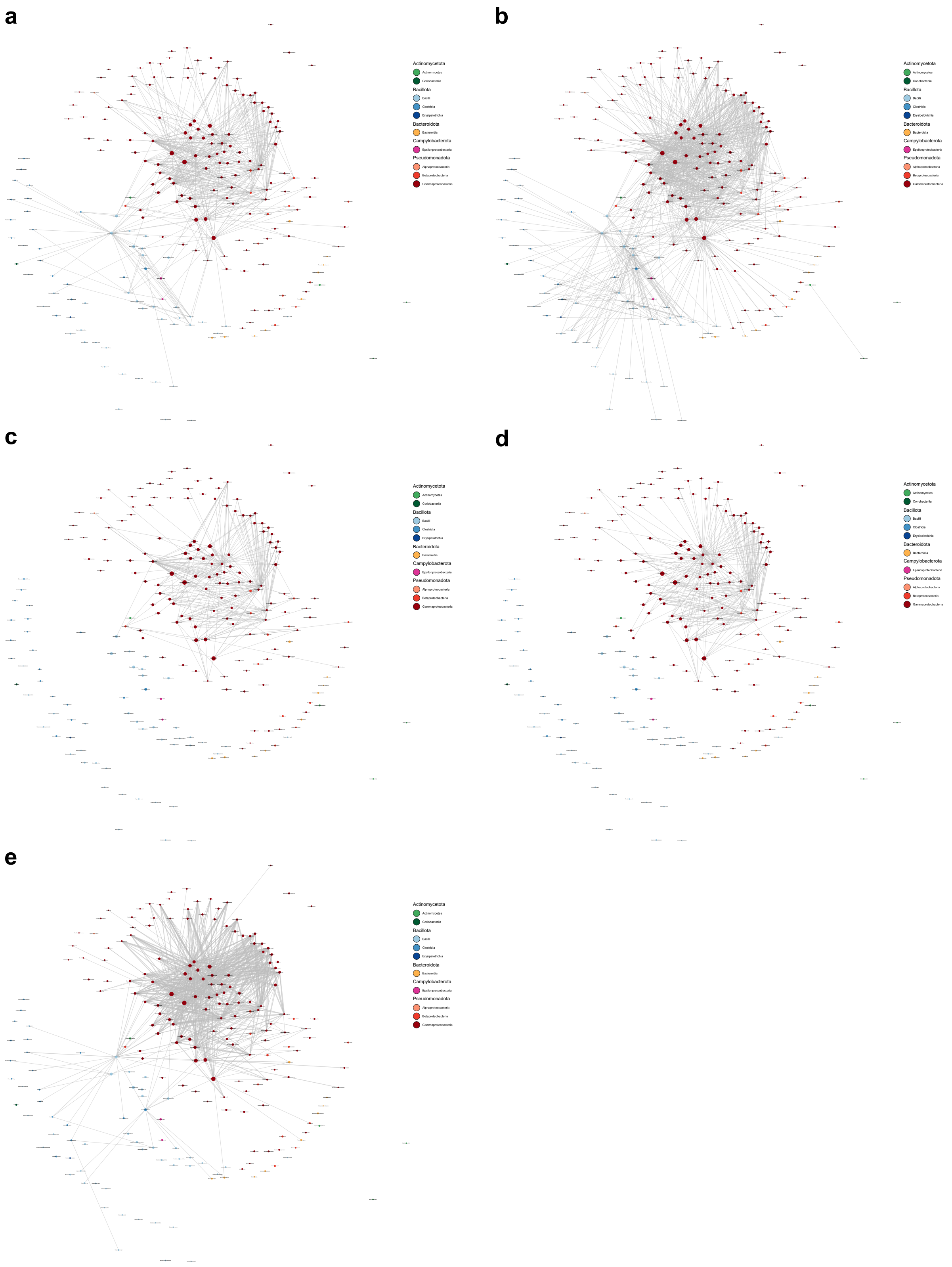


**Supplementary Fig. 8 Networks displaying the co-occurrence of promiscuous bacterial species.** Each node represents a species, the size is proportional to the total number of inferred interactions associated with that species. Edges are drawn between species with a ≥ order-level distance, between which horizontal transfer was observed ≥ 5 times. Edge thickness indicates the maximal estimated co-occurrence of two species included in each taxon in **a** Animal samples (n=4,376), **b** Human samples (n=3,220), **c** Soil samples (n=4,137), **d** Water samples (n=7,898), and **e** Wastewater samples (n=1,185). If co-occurrence was measurable in less than 1% of the corresponding samples, no edge is drawn.

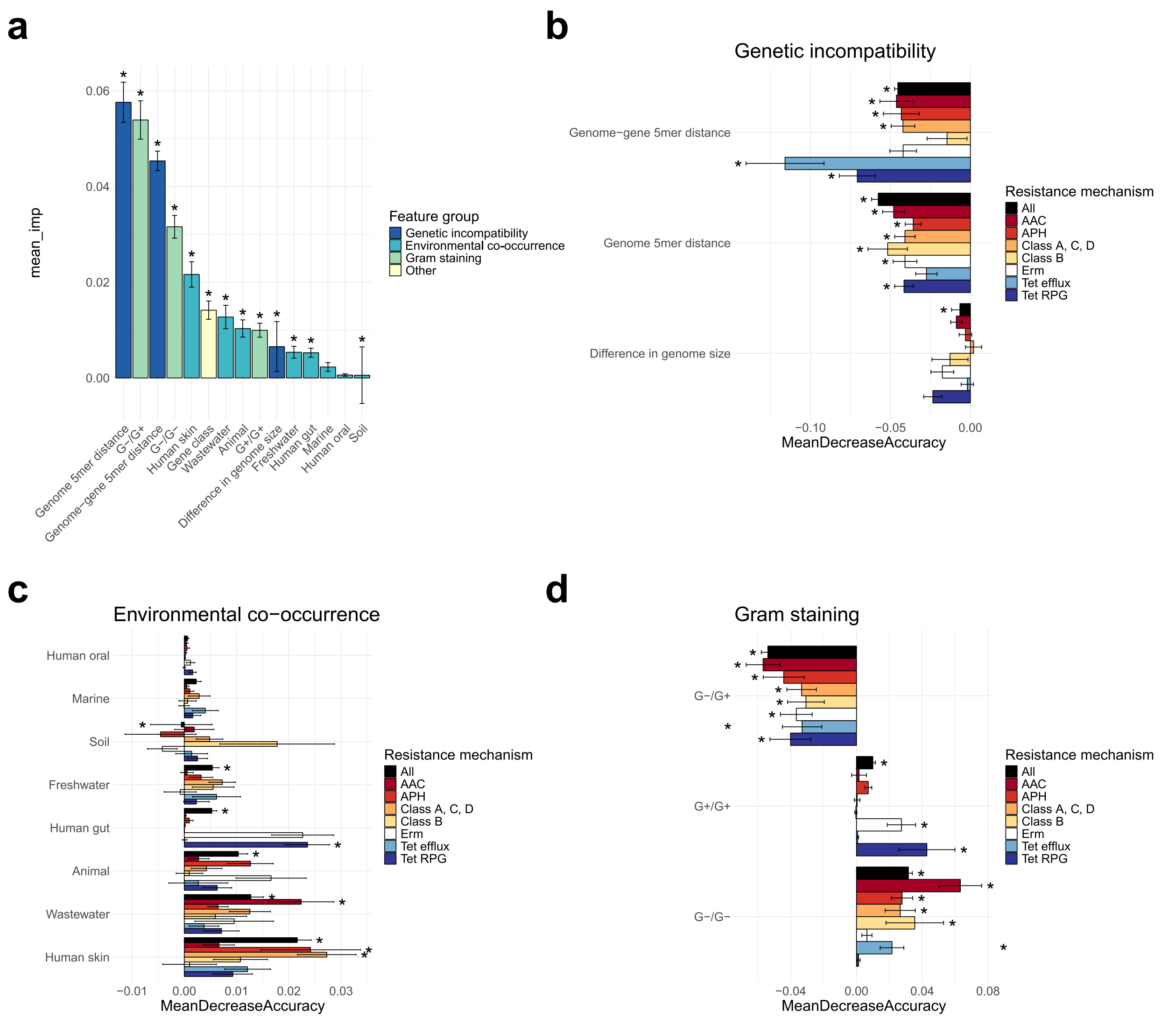


**Fig. 3. Relative importance of factors from models using more specific co-occurrence variables. a** The importance of the factors included in the general random forest classifier over ten iterations, ordered according to their overall contribution to the accuracy of the classifier (MeanDecreaseAccuracy). Standard deviations are included as error bars, and factors that were significant (*p*-value<0.01) across all ten iterations are denoted by an asterisk. **b**–**d** The mean importance of each factor group (genetic incompatibility, environmental co-occurrence, and Gram staining, respectively) for all eight random forest classifiers over ten iterations. Signs have been added to show whether an increased value of the variable is generally indicative of horizontally spread ARGs (+) or not (–).

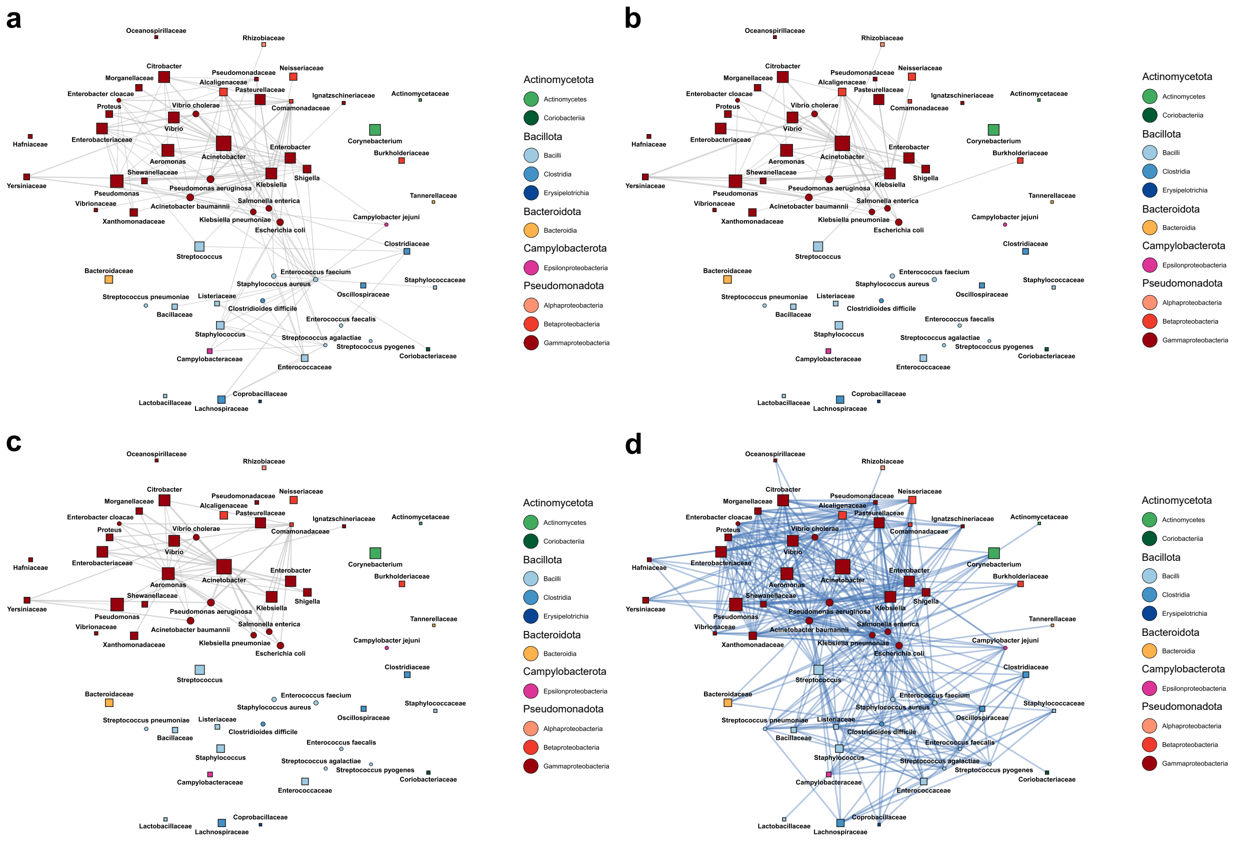


**Supplementary Fig. 10 Networks displaying the co-occurrence of promiscuous bacterial taxa.** Each node represents a taxon on either species, genus, or family level, which was aggregated such that individual nodes on a lower level are not part of the corresponding higher-level node(s). For each node, the size is proportional to the total number of inferred interactions associated with that taxon, and the shape indicates if the node represents a species (circle) or a higher-level taxon (square). Edges are drawn between taxa with a ≥ order-level distance, between which horizontal transfer was observed ≥ 5 times. Edge thickness indicates the maximal estimated co-occurrence of two species included in each taxon in **a** Animal samples (n=4,376), **b** Soil samples (n=4,137), and **c** Water samples (n=7,898). If co-occurrence was measurable in <1% of the corresponding samples, no edge is drawn. In panel **d** no co-occurrence is represented, instead all frequently observed interactions (≥ 5 transfers) are included as edges of equal thickness.

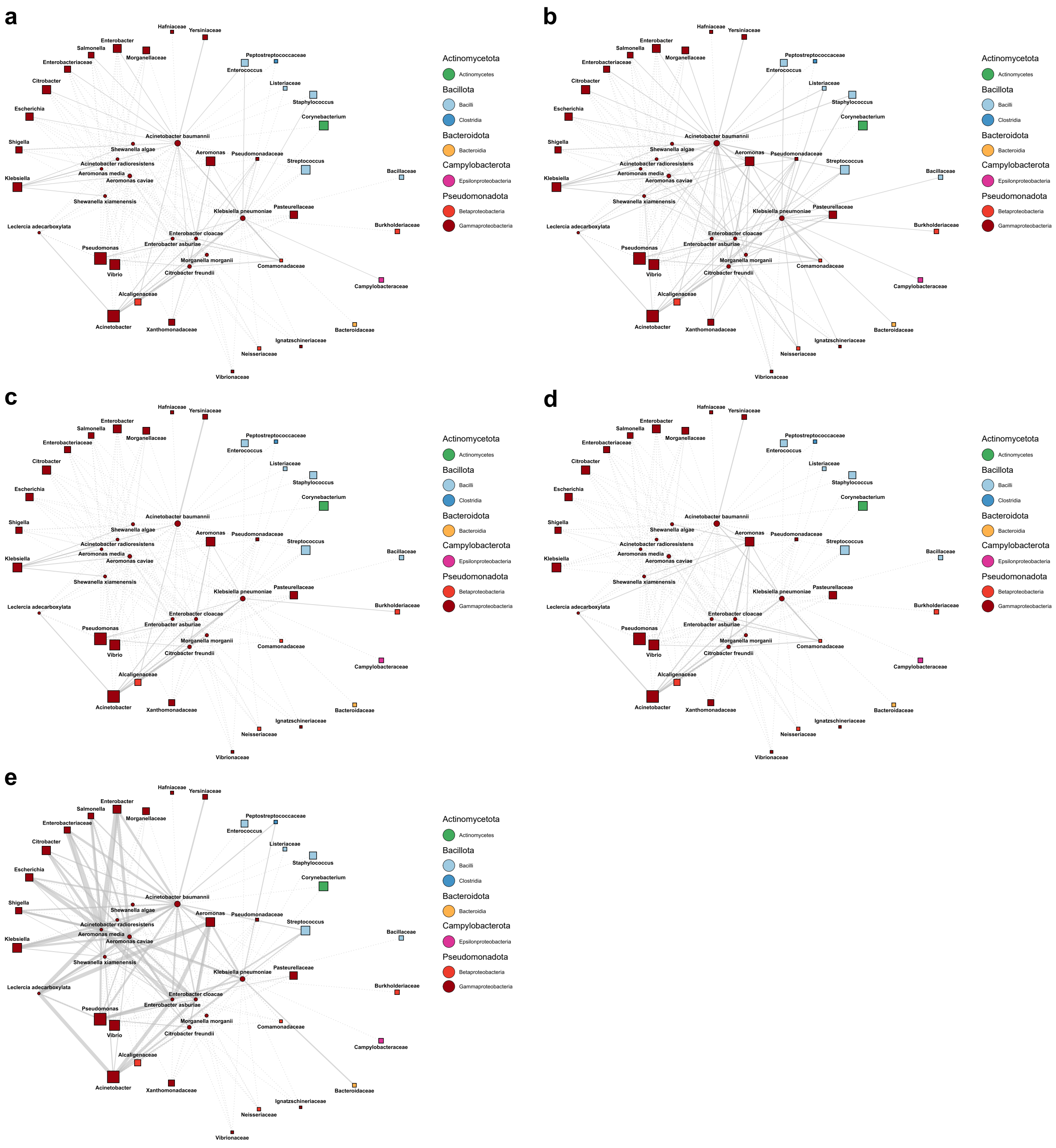


**Supplementary Fig. 11 Networks displaying the observed gene transfers and co-occurrence between origin species and other bacterial taxa.** Each node represents either an origin species or a taxon on either genus, or family level, which was aggregated such that individual nodes on a lower level are not part of the corresponding higher-level node(s). For each node, the size is proportional to the total number of inferred interactions associated with that taxon, and the shape indicates if the node represents an origin species (circle) or a higher-level taxon (square). Edges are only drawn between origin species and other taxa with a ≥ order-level distance, between which horizontal transfer was observed ≥ 5 times. Edge thickness indicates the maximal estimated co-occurrence of two species included in each taxon in in **a** Animal samples (n=4,376), **b** Human samples (n=3,220), **c** Soil samples (n=4,137), **d** Water samples (n=7,898), and **e** Wastewater samples (n=1,185). If co-occurrence was measurable in less than 1% of the corresponding samples, the connection is represented as a dashed line.

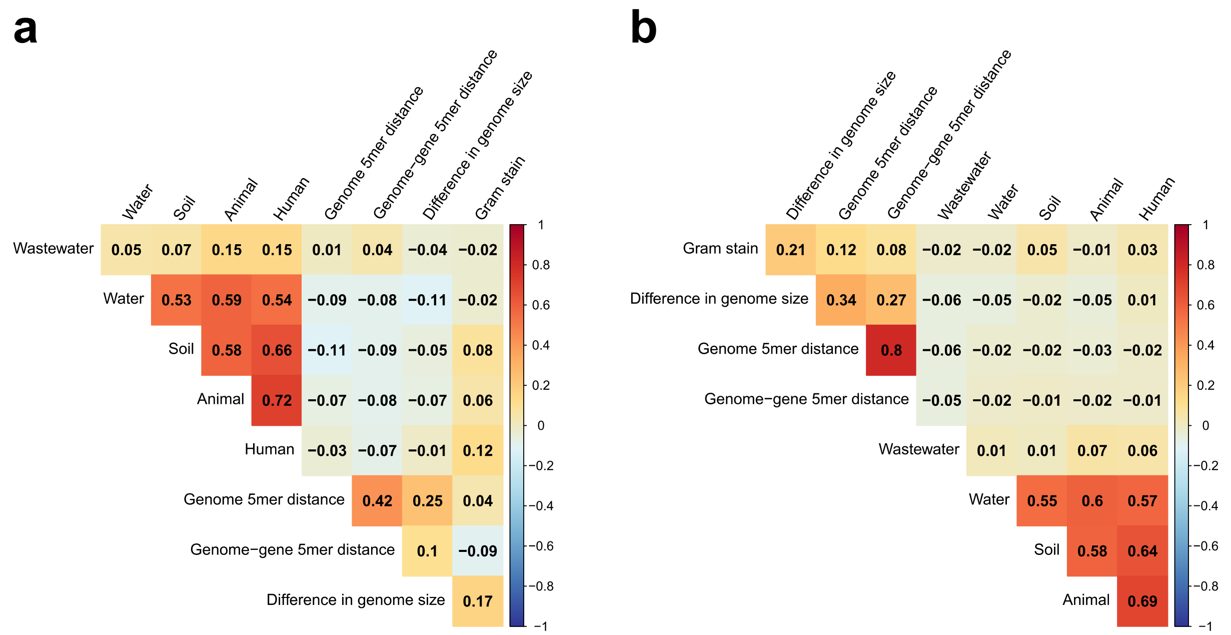


**Supplementary Fig. 12 Spearman's correlation of the features used to train random forest classifiers. a** Observed transfers. **b** Randomized transfers.

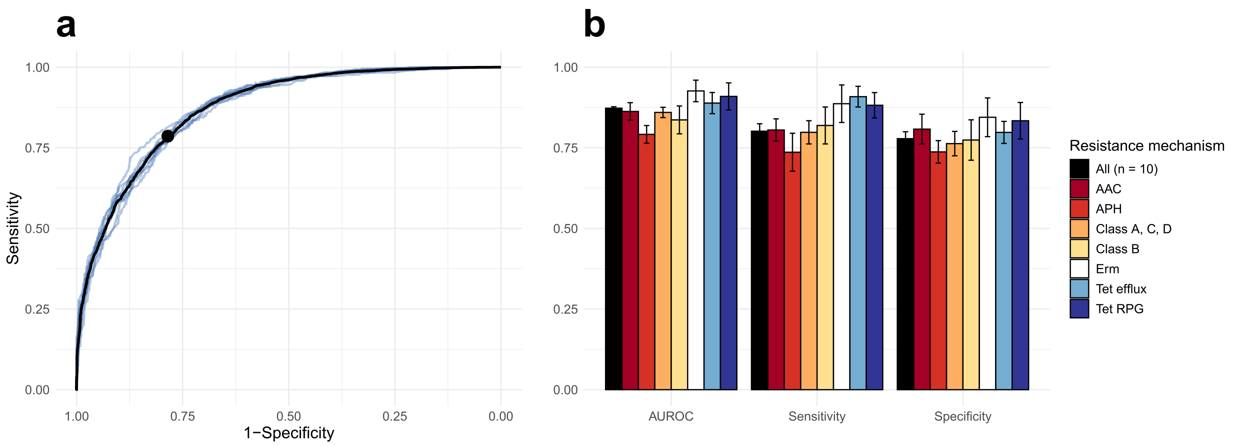


**Supplementary Fig. 13. Performance of models using more specific co-occurrence variables. a** Receiver operating characteristic curves produced from predictions on test data by random forest models trained on horizontal transfers representing all included resistance mechanisms, over ten iterations. Each model was built using variables representing the genetic incompatibility, environmental co-occurrence, and cell wall composition of the bacteria involved in each transfer. The black line represents the mean of the produced receiver operating characteristics (ROC) curves. The point represents the mean optimal performance (the point closest to a sensitivity and specificity of 1). **b** Area under the ROC curve (AUROC), sensitivity, and specificity observed for predictions on test data using random forest models representing different resistance mechanisms with enough data present (>100 transfers observed). Each bar represents the mean of the observed metric over ten iterations, with error bars representing the standard deviation.

**Supplementary Table 1. Summary of predicted ARGs and horizontally spread ARGs**

| Gene type | Predicted ARGs [unique protein sequences] | Detected horizontal transfers [unique points in the tree] |
|---|---|---|
| **Aminoglycoside** | | |
| AAC(2’) | 3,474 [879] | 24 [24] |
| AAC(3) class 1 | 2,467 [295] | 62 [46] |
| AAC(3) class 2 | 114,495 [3,853] | 163 [98] |
| AAC(6’) class 1 | 22,520 [1,081] | 258 [83] |
| AAC(6’) class 2 | 471,068 [2,512] | 69 [58] |
| AAC(6’) class 3 | 22,164 [227] | 31 [17] |
| APH(2’’) | 3,729 [568] | 33 [24] |
| APH(3’) | 252,441 [3,368] | 1,198 [241] |
| APH(6) | 198,264 [5,010] | 648 [186] |
| **Beta-lactam** | | |
| Class A | 291,085 [12,399] | 1,196 [353] |
| Class B1/B2 | 12,713 [2,017] | 130 [85] |
| Class B3 | 461,461 [3,255] | 70 [70] |
| Class C | 90,543 [7,329] | 60 [53] |
| Class D1 | 40,392 [1,536] | 73 [27] |
| Class D2 | 126,394 [3,274] | 163 [89] |
| **Macrolide** | | |
| Erm type A | 1,433 [461] | 53 [39] |
| Erm type F | 56,706 [938] | 310 [109] |
| Mph | 62,924 [1,551] | 140 [52] |
| **Quinolone** | | |
| Qnr | 53,980 [1,670] | 133 [27] |
| **Tetracycline** | | |
| Tet efflux | 262,452 [3,161] | 700 [103] |
| Tet enzyme | 2,212 [448] | 57 [36] |
| Tet RPG | 113,085 [4,964] | 705 [324] |
| **Total** | **2,666,002 [60,773]** | **6,276 [2,144]** |
